# Potential of Photoelectric Stimulation with Ultrasmall Carbon Electrode on Neural Tissue: New Directions in Neuromodulation Technology Development

**DOI:** 10.1101/2024.02.17.580823

**Authors:** Keying Chen, Bingchen Wu, Daniela Krahe, Alberto Vazquez, James R. Siegenthaler, Robert Rechenberg, Wen Li, X. Tracy Cui, Takashi D.Y. Kozai

## Abstract

**Objective:** Neuromodulation technologies have gained considerable attention for its clinical potential in treating neurological disorders and their capacity to advance cognition research. Nevertheless, traditional neuromodulation methods such as electrical stimulation and optogenetics manipulation currently experience technical and biological challenges that hinge their therapeutic potential and chronic research applications. Recently, a promising alternative neuromodulation approach based on the photoelectric effect has emerged. This approach is capable of generating electrical pulses when exposed to near-infrared (NIR) light and allows modulation of neuronal activity without the need for genetic alterations. In this study, we investigate a variety of design strategies aimed at enhancing photoelectric stimulation using minimally invasive, ultrasmall, untethered carbon electrodes.

**Approach:** A multiphoton laser was employed as the NIR light source. Benchtop investigations were conducted using a three-electrode setup, and chronopotentiometry was used to record photo-stimulated voltage. For *in vivo* evaluation, we used Thy1-GCaMP6s mice with acute implantation of ultrasmall carbon electrodes.

**Main results:** We revealed the beneficial effects of high duty-cycle laser scanning and photovoltaic polymer interfaces on the photo-stimulated voltages of ultrasmall carbon electrodes. Additionally, we demonstrated the promising potential of carbon-based diamond electrodes for photoelectric stimulation and examined the application of photoelectric stimulation in precise chemical delivery by loading mesoporous silica nanoparticles (SNPs) co-deposited with polyethylenedioxythiophene (PEDOT).

**Significance:** These findings on photoelectric stimulation utilizing ultrasmall carbon electrodes underscore its immense potential for advancing the next generation of neuromodulation technology. This approach offers the opportunity to effectively modulate neural tissue while minimizing invasive implantation-related injuries in freely moving subjects, which hold significant promise for a wide range of applications in neuroscience research and clinical settings.

## 1. Introduction

The neuromodulation technology field is experiencing rapid growth, capturing the interest of both the scientific community and the general public due to its potential in treating a variety of neurological and neuropsychiatric conditions [1–7]. Neuromodulation involves directly interfacing with the nervous system using electrical, electromagnetic, or chemical methods to precisely control and regulate neural activity [8–10]. These neuromodulation technologies stand out from pharmaceuticals and surgical interventions by offering greater spatial and temporal precision as well as the benefit of being reversible [2]. Demonstrated successes in managing diseases like Parkinson’s, depression, epilepsy, and spinal cord injuries have fueled a push to improve current technologies and develop new approaches to expand the scope of neuromodulation for wider clinical use [2, 6, 8, 11].

For decades, neuromodulation has primarily been accomplished through traditional electrical stimulation, now a common technique in clinical settings [12–14]. Electrically controlled drug delivery with implantable electrode arrays has attracted recent attention because of its ability to deliver neuromodulation with spatial precision. Localized drug delivery enables targeted therapy to the region of interest in the neural tissue and enhances the effectiveness of the treatment without the inherent side effects of systemic treatments [15, 16]. Fluidic delivery methods using microcapillary or microfluidic channels have been traditionally used to deliver payloads to the neural tissue. These techniques are prone to failures such as leakage, clogging, and reflux. On top of that, these systems require additional hardware, such as pumps and valves that make them complex and usually bulky. Moreover, incorporating fluidic channels into neural devices increases the footprint of the devices, rendering them more invasive. Also, the pressurized delivery method might cause local edema [17–23]. Electrically controlled drug delivery does not require solvent assistance; thus this “dry” delivery technique could deliver drugs without disrupting local pressure.

More recently, optogenetics has also risen in prominence for its temporal precision in controlling neural activity with visible light and its capacity to dissect neural circuitry by targeting specific neuron subtypes [10, 24, 25]. Yet, modulating deeper brain structures typically requires invasive implantation of electrodes and optical fibers due to the screening of electric fields and scattering of light within brain tissue [26, 27]. Chronic implantation with large, invasive devices can result in a foreign body response, leading to glial scar formation and neuronal degeneration [28–31]. Moreover, tethering animals with electrode wires or light sources during behavioral studies can prevent subjects from freely moving and increase the risk of brain implant displacement, infection, and mechanical failure [32, 33]. These technical and biological challenges are factors affecting both the therapeutic potential for treating neurological disorders and the research potential for longitudinal modulation studies.

Several improvement methods have been reported to mitigate these challenges in neuromodulation. Reducing implant sizes to the sub-cellular scale, using flexible materials in device construction, and developing wireless transmission and packaging techniques have all contributed to improving tissue compatibility, reducing mechanical stress, and enhancing long-term biocompatibility [34–41]. Additionally, the development of wireless optogenetic interfaces and new opsins responsive to near-infrared (NIR) light has enabled the modulation of deep subcortical structures, thereby increasing the spatial precision of light-based neuromodulation [42–44]. However, there remain limited studies on neuromodulation devices that can simultaneously eliminate the need for head tethering, minimize brain injury associated with implantation, and utilize light for neural stimulation. Our previous work validates an innovative strategy for neuromodulation based on the photoelectric effect, employing ultrasmall, free-floating, glass-insulated carbon fiber electrodes activated by NIR multiphoton laser stimulation [45]. This method enables a less invasive, untethered approach to stimulate neurons effectively, without the need for genetic alterations typically used in optogenetics.

The concept of stimulation based on the photoelectric effect originates from the voltage artifacts generated during simultaneous multiphoton imaging and electrophysiological recording with metallic electrodes [46]. These constantly occurring photoelectric artifacts are challenging to eliminate but present an opportunity to be leveraged for beneficial purposes. Electrodes made from doped semiconductor materials, like carbon fibers, can produce a bulk photovoltaic voltage due to the series arrangement of grains or domains [46, 47]. When activated by NIR multiphoton laser stimulation, these carbon electrodes induce each domain to generate a small electron flow for voltage. These small voltages cumulatively result in a transient, non-Faradaic, capacitive charge transfer at the electrode-electrolyte interface upon switching the light source on or off. *In vivo* data has demonstrated that such multiphoton laser stimulation photoelectrically activates neuronal activity in Thy1-GCaMP mice, with minimal thermal heat generated [45]. However, the current model with glass-insulated carbon fiber electrodes has limitations in flexibility and intensity of the photoelectric effect. Therefore, there is a need to develop strategies that improve photoelectric stimulation techniques for modulating neural activity in the brain.

In this work, we tested and validated several different strategies to improve the photoelectric stimulation with ultrasmall carbon electrodes. We first developed an electrode prototype involving a free-floating 7 μm diameter carbon fiber electrode with flexible, transparent parylene-C insulation. This design minimizes the risk of tissue damage, but the parylene insulation is prone to damage unless stimulated with low laser power. Then we found that a higher duty-cycle laser scanning increased the photo-stimulated voltage. We also investigated the effects of a special organic photovoltaic polymer blend of poly(3-hexylthiophene) (P3HT) and [6,6]-phenyl C61-butyric acid methylester (PCBM) at the electrode interfaces. With a strong photovoltaic effect, this P3HT: PCBM photovoltaic polymer blends are widely used in solar cells and recently have applications in wafer-based *in vitro* biological stimulation [48–53]. This polymer coating promotes neuronal activation with lower laser power to avoid phototoxic damage and maintain the integrity of parylene insulation. Furthermore, we validated the efficacy of an alternative carbon-based electrode -- the ultrasmall diamond electrode -- for neuronal activation *in vivo*. Lastly, we revealed the potential of photoelectric stimulation technology in on-demand chemical delivery with high spatial and temporal resolution. Overall, these results demonstrate that photoelectric stimulation with ultrasmall carbon electrodes is an effective neuromodulation approach worth further investment in future research, which has the potential to broaden the scope of its application in neuroscience research and clinical settings.

## 2. Methods

### 2.1. Ultrasmall carbon electrode preparation

#### 2.1.1 Parylene-insulated carbon fiber electrode

Electrodes were prepared using a single 7-micron diameter carbon fiber (Cytec Thornel T650) (**Fig. 1A**). The single carbon fiber was threaded into borosilicate glass capillaries (0.4 mm ID, 0.6 mm OD; A-M systems Inc.) that was pulled to a fine glass tip using a vertical electrode puller (Narishige puller). To maintain its position securely within the glass tip, the carbon fiber was sealed with low-viscosity epoxy (Spurr Epoxy; Polysciences Inc) and left to dry overnight. Electrodes with photovoltaic coating were involved in additional procedures for polymer deposition. The carbon fiber was first cleaned by isopropyl alcohol soaking. Poly(3-hexylthiophene) (P3HT, regio-regularity of 99.5 %, Millipore Sigma) and [6,6]-Phenyl C61 butyric acid methyl ester (PCBM, Sigma-Aldrich) were diluted to 45 mg/mL and mixed (1:1 volume ratio) using a magnetic stirrer. The mixture solution was then heated to 50 degrees Celsius for 20 min and deposited onto the carbon fiber by manually dip coating. Coated fibers were cured at 120 degrees Celsius for 10 min. Subsequently a 2 μm layer of Parylene-C was deposited (di-chloro-di-p-xylylene) as insulation. The exposed tip (∼ 100 µm) was sharpened by flame-burning as elucidated in a previous study [54], where the remaining fiber was submerged in water to allow controlled burning of the protruding tips down to the water’s surface level (**Fig. 1B**).

**Figure 1.**
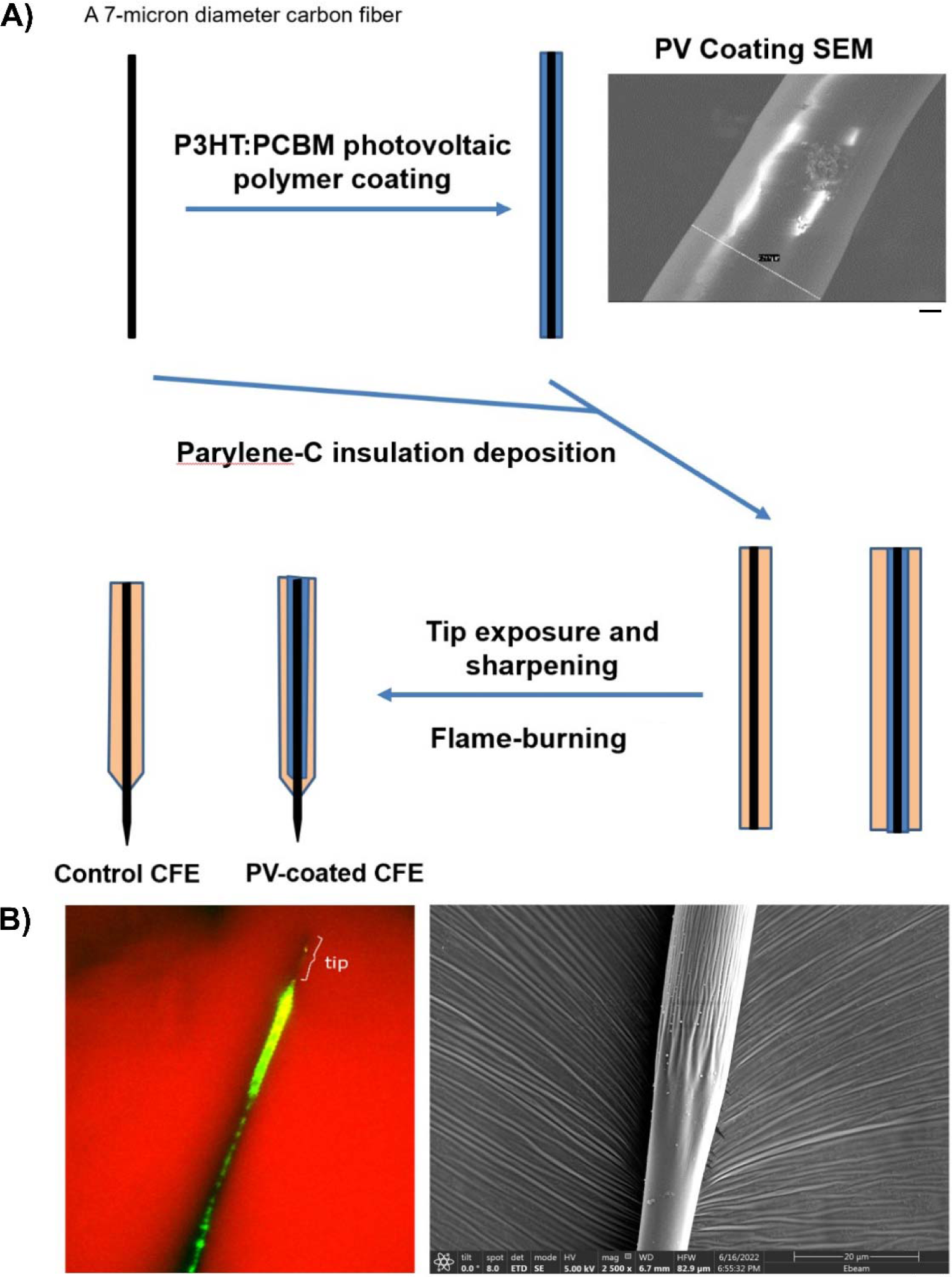
Prototype electrode preparation. A) Schematic diagram of prototype electrode preparation. Representative scanning electron microscopy (SEM) image of P3HT:PCBM polymer coating uniformly deposited onto carbon fiber. The control carbon fibers without photovoltaic (PV) coating skipped this step. Scale bar = 1 μm. Then, the carbon fiber tip was cleaned by isopropyl alcohol and deposited by 2 μm layer of Parylene-C (di-chloro-di-p-xylylene). Finally, the tip was exposed and sharpened by flame-burning. The remaining array was held underwater so that the protruding fiber tip was burned down to the surface of the water B) Left: Representative two-photon image of prototype electrode tip. Bare carbon fiber tip is outlined. Green segment is the parylene-C insulation region with photovoltaic polymer coating. Red background is SR101 mixed agarose. Scale bar = 50 μm. Right: Representative SEM image of prototype electrode exp Scale bar = 5 μm. sed tip.

#### 2.1.2. Diamond electrode preparation

Boron-doped polycrystalline diamond (BDD) microelectrodes were constructed similarly towards a CFME and all diamond BDD microelectrode using modifications from previously reported methods [55]. Briefly, a 4 µm BDD film was grown on a 4” Ø - 500 µm thick single-side polished silicon wafer using a 915 MHz microwave chemical vapor deposition reactor. Diborane was added during the growth at a ratio of 37,500 ppm B/C ratio to ensure conductivity. Following BDD growth, a titanium adhesion layer and a copper layer were thermally evaporated (Auto 306; Edward, INC.,West Sussex, UX). Subsequently these layers were patterned into 1 cm x 10 or 20 µm strips via photolithography (ABM-USA, INC., Jan Jose, CA, USA) and a wet chemical etching of titanium and copper with copper etchant (Millipore Sigma) and titanium with buffered hydrofluoric acid (BHF, Millipore Sigma). Diamond strips were then patterned by reactive ion etching using SF_6_/Ar/O_2_ with a microwave power of 1000 W and a radio-frequency (RF) bias of 150 W (180 V). Finally, the diamond strips were released from the silicon using an HNA etchant which is a mixture of hydrofluoric, nitric, acetic.

The boron-doped diamond microelectrodes (BDDMEs) were fabricated by first attaching a BDD strip onto a 32 AWG wire wrapping wire using EPO-TEK H20E (Epoxy Technology, Billerica, MA, USA) conductive silver epoxy. The epoxy was heat cured for 30 min at 110 ⁰C. The wire and BDD were then threaded into a pulled glass capillary with an open tapered tip (World Precision Instruments. Sarasota, FL, USA), and sealed in place using an insulating 2-part, 5-min epoxy (J-B Weld, Marietta GA, USA), and cured overnight. Electrodes were pretested using fast-scan cyclic voltammetry (FSCV) to ensure functionality and stability. The electrode underwent potential sweeping between 1.3 V and -0.4 V at a scan rate of 400 V/s, repeated at 10 Hz until the electrode stabilized in Tris aCSF.

#### 2.1.3. PEDOT/SNP coated carbon fiber electrode preparation

Mesoporous, sulfonate-modified silica nanoparticles (SNPs) were synthesized via a previously reported method [56]. Hexadecyl trimethylammonium bromide was first added to a mixture of triethanolamine, water, and ethanol to form the nanotubes which eventually would constitute the pores. Next, tetraethyl orthosilicate and mercaptopropyl-trimethoxysilane were added to form the thiolated silica nanoparticles around said pores. Surfactant within the pores was removed with an acid wash, and thiol groups were converted to sulfonate groups by the addition of hydrogen peroxide solution; this additional step allows the nanoparticles to be used as a conducting dopant. Polymerizations were carried out with a three-electrode setup with a platinum foil counter, carbon fiber working electrode, and Ag/AgCl reference. Ethylenedioxythiophene (EDOT, 0.01 Mol) in deionized (DI) water was mixed with SNP (5 mg/mL) and sonicated for 5 min in a bath sonicater. Polymerization is carried out under constant current (320 µA/cm^2^) to 160 mC/cm^2^. For loading SNP, fluorescein (25 mg/mL) was dissolved in water and 5 mg of mesoporous SNP was sonicated with 200 µL of fluorescein solution for 20 min prior to the particles being collected by centrifuge and dried under vacuum.

### 2.2. Benchtop evaluation to measure photo-stimulation induced voltages

A three-electrode setup was used to evaluate the voltage generated by the carbon electrode during photo-stimulation [45]. The phantom used in this setup was prepared using 50 mL of saline mixed with 0.5% agar gel. Sulforhodamine 101 (1 μMol) for background visualization was added to the agar mixture after heating. A bundle of carbon fibers served as the counter electrode (Cytec Thornel T650), and an Ag|AgCl electrode was connected to the reference electrode. The working electrode was connected to a carbon electrode that was inserted into the phantom at a 30° angle. The photo-stimulation process was conducted utilizing a two-photon microscope from Bruker (Middleton, WI) equipped with an ultra-fast laser (Insight DS+; Spectra-Physics, Menlo Park, CA) tuned to an 800 nm wavelength and a 4.0 µs dwell time. The system featured non-descanned photomultiplier tubes provided by Hamamatsu Photonics KK (Hamamatsu, Shizuoka, Japan) and employed a 16X, 0.8 numerical aperture water immersion objective lens sourced from Nikon Instruments (Melville, NY). The point spread function of our two-photon microscopy system is ∼ 7 μm in the z-direction. Attention was devoted to maintaining the laser focus on the prototype electrodes during photos-stimulation.

Chronopotentiometry was performed using an Autolab unit with an ECD (extremely low current) module and NOVA software (PGSTAT128N, Metrohm Autolab, Utrecht, Netherlands) to measure the generated voltage by the carbon electrode. For the Electrochemical impedance spectroscopy (EIS) measurements, impedances were recorded from the carbon electrode connected to the Autolab potentiostat. Impedances were recorded for each channel using a 10 mV RMS sine wave in a range of 1 Hz–100 kHz. For the cyclic voltammetry (CV) tests, the working electrode potential was swept between 0.8 V and −0.6 V with a scan rate of 1 V/s.

### 2.3. Electrode implantation surgery

The carbon electrode was implanted in Thy1-GCaMP6s transgenic mice (N = 3) for *in vivo* evaluation. Initially, the mice were anesthetized using an intraperitoneal (IP) injection comprising 75 mg/kg of ketamine and 7 mg/kg of xylazine [57]. Supplementary doses of ketamine (40 mg/kg) were administered to maintain anesthesia throughout the surgery. Mice were head-fixed onto a stereotaxic frame on a heating pad and administered with a continuous supply of oxygen (1 L/min). A craniotomy window (3 × 3 mm) was prepared over the left visual cortex in coordinates of 1 mm anterior to lambda and 1.5 mm lateral from the midline. To ensure brain hydration and prevent thermal damage, saline was periodically applied during the procedure. The carbon electrode was then inserted 400 μm into the cortex at a 30° angle (insertion velocity of 200 μm/s, oil hydraulic Microdrive; MO-82, Narishige, Japan). This insertion avoided large vessels and positioned the tip approximately 200 µm below the brain surface. All procedures and experimental protocols were approved by the University of Pittsburgh, Division of Laboratory Animal Resources and Institutional Animal Care and Use Committee in accordance with the standards for humane animal care as set by the Animal Welfare Act and the National Institutes of Health Guide for the Care and Use of Laboratory Animals.

### 2.4. *In vivo* neuronal calcium imaging

Photo-stimulation utilized the two-photon laser employing two methods: 0.2 Hz raster scanning across the electrode or 1-second (1-s) repetitive line scanning focused near the electrode’s tip. Time-series stacks (galvo scanning, 0.69 frames per second) were captured to observe neuronal calcium responses around the implanted prototype electrodes. To avoid laser scanning on the electrode until the photo-stimulation onset, the position of the implanted carbon electrode was masked during imaging. Imaging of the surrounding brain tissue was conducted both before and after photo-stimulation at 20 mW, 920 nm wavelength, enabling visualization of neuronal calcium activity. Neurons were deemed activated if their calcium elevation surpassed the threshold set at 3 standard deviations above the mean calcium fluorescence. The count of activated neurons was conducted in 50 μm spaced bins around the electrode. Subsequently, the activated neurons, induced by photo-stimulation, were categorized based on their temporal dynamics: 1) Onset of activation: termed “Rapid Onset” if there was an immediate increase in calcium levels within 2 seconds post photo-stimulation, and “Delayed Onset” if the elevation occurred 15 seconds or later post photo-stimulation. 2) Duration of activation: classified as “Sustained” if activated neurons maintained elevated calcium levels for over 30 seconds during the post-stimulation period, and “Transient” if the calcium increase persisted for less than 10 seconds.

### 2.5. Data Analysis

Data analysis was performed in MATLAB (Mathworks, Natick, MA). Neurons activated with increased calcium activity near the prototype electrode were manually identified. Changes in calcium fluorescence (dF) were normalized to the mean intensity of the pre-stimulation period (F_o_). Data was averaged across all animals and plotted as mean ± standard error.

### 2.6. Statistics

One-way ANOVA with Tukey post hoc (*p* < 0.05) was used to determine any significant difference in photo-stimulated voltage across varying laser powers. Welch’s unpaired t-test was used to discern significant differences in generated voltage between laser scanning methods. Two-way ANOVA with Bonferroni corrected t-test (*p* < 0.05) was applied to find significant differences between photovoltaic coated prototype electrodes and uncoated controls in generated voltage at various laser power and the number of activated neurons at distance bins.

## 3. Results

Photoelectric stimulating carbon electrodes with infrared light has emerged as a promising strategy for brain tissue stimulation. A previous study has demonstrated that carbon fiber electrodes with glass insulation can generate voltage pulses during photo-stimulation, effectively activating neurons close to the electrode site [45]. To advance this technology, our aim was to test several design strategies and assess their potential to improve photoelectric stimulation with carbon electrodes: 1) Implementing flexible, transparent Parylene C insulation; 2) Exploring different light source scanning methods; 3) Incorporating photovoltaic polymer interfaces; 4) Evaluating the use of alternative carbon-based diamond electrodes; 5) Investigating the integration of a conductive polymer drug delivery system. The results of this study would help establish novel directions to illuminate the future development of photoelectric stimulation technology for neuromodulation.

### 3.1. Flexible Parylene C insulation is easily damaged at high stimulating laser intensity

A prior study has demonstrated that a stiff glass-insulated carbon fiber electrode can effectively generate photo-stimulated voltage for *in vivo* neuronal activation, with a linear increase observed as stimulating laser intensity increases [45]. Thus, our initial improvement strategy focuses on replacing the glass insulation with a more flexible and transparent material. To implement this strategy, we designed a prototype electrode that involves a free-floating, 7 μm diameter carbon fiber electrode with a transparent and flexible parylene-C insulation. To evaluate whether our prototype electrode could successfully generate sufficient photo-stimulated voltage at different laser power intensities, we employed chronopotentiometry to measure the potential and collected two-photon time series stacks to assess the electrode integrity. Photo-stimulation was performed by rastering the two-photon scanning laser across the field of view (0.2 Hz, 800 nm wavelength, 4.0 µs dwell time), resulting in a cathodic potential generated each time the laser passed over the electrode (**Fig. 2A**). The trial-average voltage at 10 mW laser power was approximately 110 mV, and parylene insulation appeared to be intact (**Fig. 2B**).

**Figure 2.**
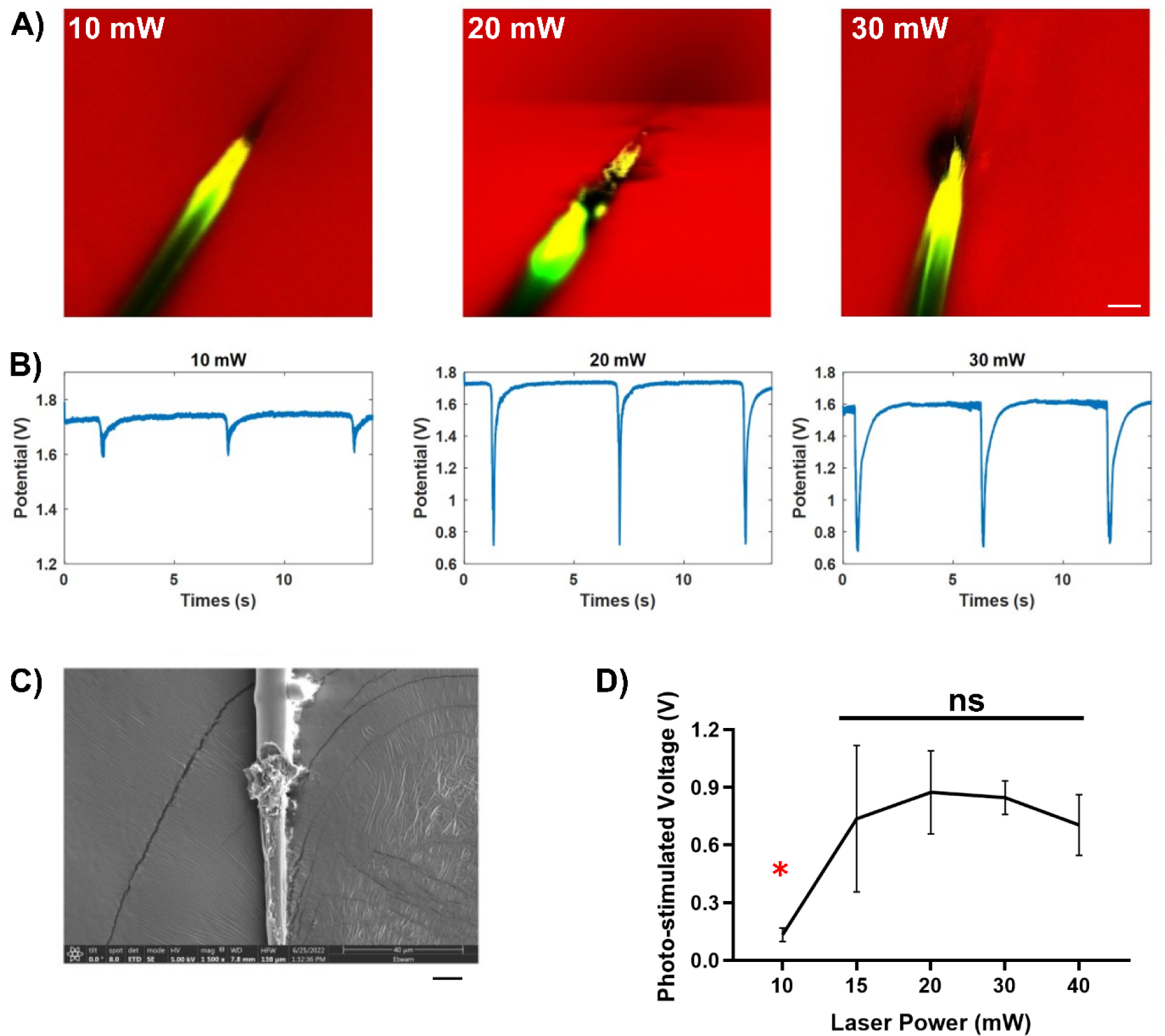
Low laser power avoids insulation damage of prototype electrode. A) Time projections of two-photon images to evaluate the stability of prototype electrodes. Parylene insulation was damaged when the laser power exceeds 10mW. B) Representative cathodic potential pulses generated by two-photon laser raster scanning at various power intensities. The trial-average voltage at 10 mW laser power was nearly 110 mV. However, the photo-stimulated voltages at 20 mW and 30 mW were similar, approximately 800 mV. C) SEM image of damaged parylene insulation after 40 mW photo-stimulation. D) Quantification of frame-scanning photostimulation voltages as a function of laser power. Scale bar = 10 μm. Red asterisk indicates statistical significance detected by One-way ANOVA with Tukey post hoc, *p* < 0.01.

This observed cathodic pulse indicated a more negatively charged environment in surrounding brain tissue, suggesting photo-stimulation activates electrons near the electrode’s surface. It’s important to note that the majority of these electrons typically return to their normal orbitals after light onset, indicating that the cathodic potential generated by the ultrasmall electrode is largely non-Faradaic, or photocapacitive [58]. Such capacitive effects are considered gentler and pose a lower risk of tissue damage [45]. While the observation of small currents and charge injections suggests that some electrons may completely leave the electrode during stimulation [45], this current leakage might be associated with the potentiostat system rather than originating directly from the electrode.

Photocapacitive stimulation is distinct from photofaradaic stimulation, such as with *pn* or *pin* junction photocells that generate current. In photofaradaic stimulation, current is generated so long as the light source illuminates the photoelectric material since electrons that are separated from the electron hole immediately enter the conduction band and are directed by an electrical gradient. Photofaradaic stimulation is much similar to traditional electrical stimulation that is typically dominated by faradaic current. Even with pseudofaradaic Iridium Oxide, there is some faradaic transfer of electrons that occur at the electrode-electrolyte interface that leads to neuronal activation. In the case of pseudofaradaic materials, the polarity of stimulation is reversed and remetalization occurs before the dissolved metal can diffuse away [58].

However, increasing the laser power beyond 10 mW led to noticeable damage to the parylene-C insulation as well as the formation of gas bubbles (**Fig. 2A-B**). Scanning Electron Microscopy (SEM) images confirmed the damage to the parylene insulation following photo-stimulation at 40 mW (**Fig. 2C**). Furthermore, increasing the laser power to 20, 30, and 40 mW yielded statistically comparable photo-stimulated voltage of approximately 800 mV, which was significantly higher than that at 10 mW (**Fig. 2D**, One-way ANOVA with Tukey post hoc, p = 0.0058). These observations suggested that the measured voltage was not linearly related to laser power intensity when parylene was damaged during photo-stimulation. This type of insulation damage was not observed with the glass insulation used in previous studies [59]. Together, maintaining a lower laser intensity during photo-stimulation of the parylene-insulated electrode is necessary to prevent insulation damage and preserve the electrode function.

Regarding the unexpected insulation damage by high laser power, it becomes crucial to consider the selection and application of the appropriate insulation material for photoelectric devices. Parylene C has high flexibility, biocompatibility, and corrosion resistance, which serve as an excellent choice for constructing medical devices such as implant encapsulation of stents and pacemakers [60, 61]. However, our results (**Fig. 2**) indicate that high laser power stimulation can generate gas bubbles, potentially damaging the parylene insulation. It is uncertain whether these bubbles form underneath the parylene layer. One possible explanation could be that the focal energy by NIR stimulating laser disrupts the triple junction - the intersection of saline, parylene C insulation, and carbon fiber core. This triple gap junction is a line defect of three boundaries and is theoretically prone to fractures, which can compromise the mechanical strength of polycrystalline materials [62, 63]. The cathodic discharge induced by NIR stimulation is likely localized at the dielectric-carbon-fluid interfaces and may expand over time, leading to subsequent electrolysis of water in the agarose gel and consequent mechanical breakdown of the parylene insulation. Further research is required to unravel these mechanisms, which will aid in refining the selection and application of suitable insulation materials.

### 3.2. Alternative High Duty-Cycle Laser Stimulation at Low Power

While the 10 mW laser rastering stimulation appeared to generate limited voltage for effective neuronal stimulation, we explored alternative laser stimulation strategies to enhance the efficacy of the parylene-insulated prototype electrode. Recognizing that the 0.2 Hz raster scanning offered a limited duty cycle (with the prototype electrode being scanned every 5.5 seconds), we turned to a repetitive scanning method that focused on a line perpendicular to the electrode over the stimulation period. This approach allowed for a near 100% duty cycle when the laser was concentrated on the electrode, resulting in a sustained 1-s pulse of cathodic potential during a 1-s line scanning laser stimulation period (**Fig. 3A**). The average voltage generated by repetitive line scanning at 10 mW was 276 ± 22 mV, which was significantly higher than average voltage generated by raster scanning, 133 ± 31 mV (**Fig. 3B**, unpaired t-test, p = 0.0185). Therefore, employing laser scanning methods with a high duty-cycle could increase the photo-stimulated voltage while sustaining low laser power.

**Figure 3.**
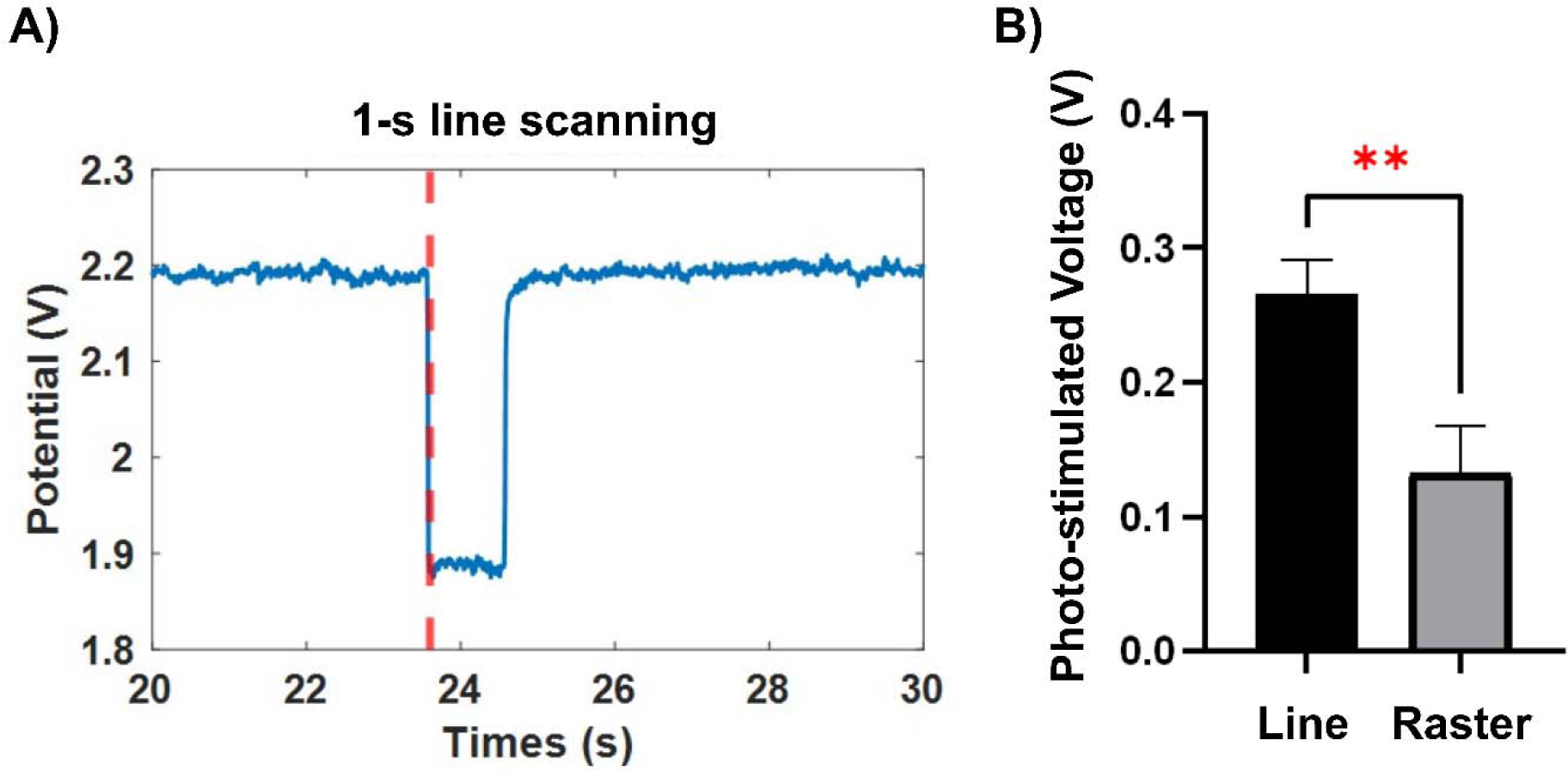
Alternative laser scanning strategy: repetitive line scanning increases photo-stimulated voltage. A) Representative cathodic potential pulse of 1-s line scanning at 10 mW photo-stimulation onto the prototype electrode. The time of laser ON is denoted as the red dash line. B) The average voltage generated by 10 mW line scanning was 276 ± 22 mV, while the average voltage by 10 mW raster scanning was 133 ± 31 mV. Red double asterisks indicate the significant difference determined by Welch’s unpaired t-test, *p* < 0.05.

The method for multiphoton laser scanning on the electrode can play a crucial role in the photoelectric-based modulation of neural activity. This method directly influences energy delivery before the light-electric conversion process. While the non-Faradic charge is generated at the onset or offset of light stimulation, the frequency of light passing the electrode is a crucial factor in determining the voltage generated during photo-stimulation [46]. This explanation may contribute to the observation that high-duty cycle stimulation as repetitive laser interactions with the electrode leads to a significantly elevated photo-stimulated voltage compared to low-duty cycle raster scanning. It is worth exploring additional laser scanning methods for future studies, as they may provide opportunities to precisely modulate the photo-stimulated voltage. One such method could involve manipulating the stimulating laser’s wavelength. Shorter wavelengths scatter more rapidly and have shallower penetration depths compared to longer wavelengths, which reduces the likelihood of photon collisions on the electrode and subsequent electron generation. While light energy decreases as wavelength increases, longer wavelengths are more likely to keep photoelectric events within safe tissue heating limits. Future studies will delve into the impact of near-infrared wavelengths to optimize the ideal wavelength for stimulation.

Our current approach places the laser scanning location near the tip of the electrodes (**Fig 2, 3**). However, it is worth noting that carbon-based electrodes are conductive and efficiently transmit energy to exposed sites along the insulated region. Initially, we hypothesized the feasibility of applying light stimulation to the base end of the electrode and the voltage would conduct to the expose electrode site. Surprisingly, this hypothesis was not realized, emphasizing that the transient voltage induced by photocapacitive stimulation results from the excitation and separation of electrons from their electron holes. The resulting voltage field was limited to the illumination site, rapidly dropping off radially without conducting to exposed regions. This explains why illuminating the insulted region of the electrode did not excite neurons, and illuminating near the edge of the insulated fiber led to insulation breakdown. Consequently, insulation plays a lesser role in shaping the voltage profile in the tissue compared to conventional electrical stimulation. Instead, the stimulation of local neuronal populations hinges on directing coherent photon beams, particularly multiphoton beams, which will play a more significant role in concentrating the stimulation on the uninsulated region of the photovoltaic electrode. Future research aimed at deep brain targets should focus on building photovoltaic electrodes coupled to waveguides rather than insulated electrical conductors.

### 3.3. Photovoltaic Polymer Coating to Increase Photo-stimulated Voltages

While employing a laser scanning strategy with a high duty cycle increased the voltage, this enhancement was not substantial. Thus, we asked whether there are additional engineering approaches to amplify the photo-stimulated voltage generated by the parylene-insulated prototype electrode. Mixtures of poly(3-hexylthiophene) and [6,6]-Phenyl C61 butyric acid methyl ester (P3HT: PCBM) have wide applications in solar cells [48, 49], with recent studies showing that this organic photovoltaic blend is effective in photo-stimulating neurons grown at the interface [51, 53]. Therefore, we applied a layer of photovoltaic (PV) polymer coating interfaced at the carbon fiber in the prototype electrode to increase the light conversion to voltage and maintain stable parylene insulation during photo-stimulation. We first examined the electrochemical properties of the PV coated prototype electrode. The results of cyclic voltammetry showed that the PV coated electrode increased the charge storage capacity compared to uncoated controls (**Fig. 4A**). Additionally, PV coated electrode exhibited a lower impedance compared to controls, especially over the low frequencies (**Fig. 4B**). Thus, PV coating helps improve electrochemical properties of carbon fiber prototype electrode, increasing charge storage capacity and decreasing the impedance over the frequency range of 1-100 kHz.

**Figure 4.**
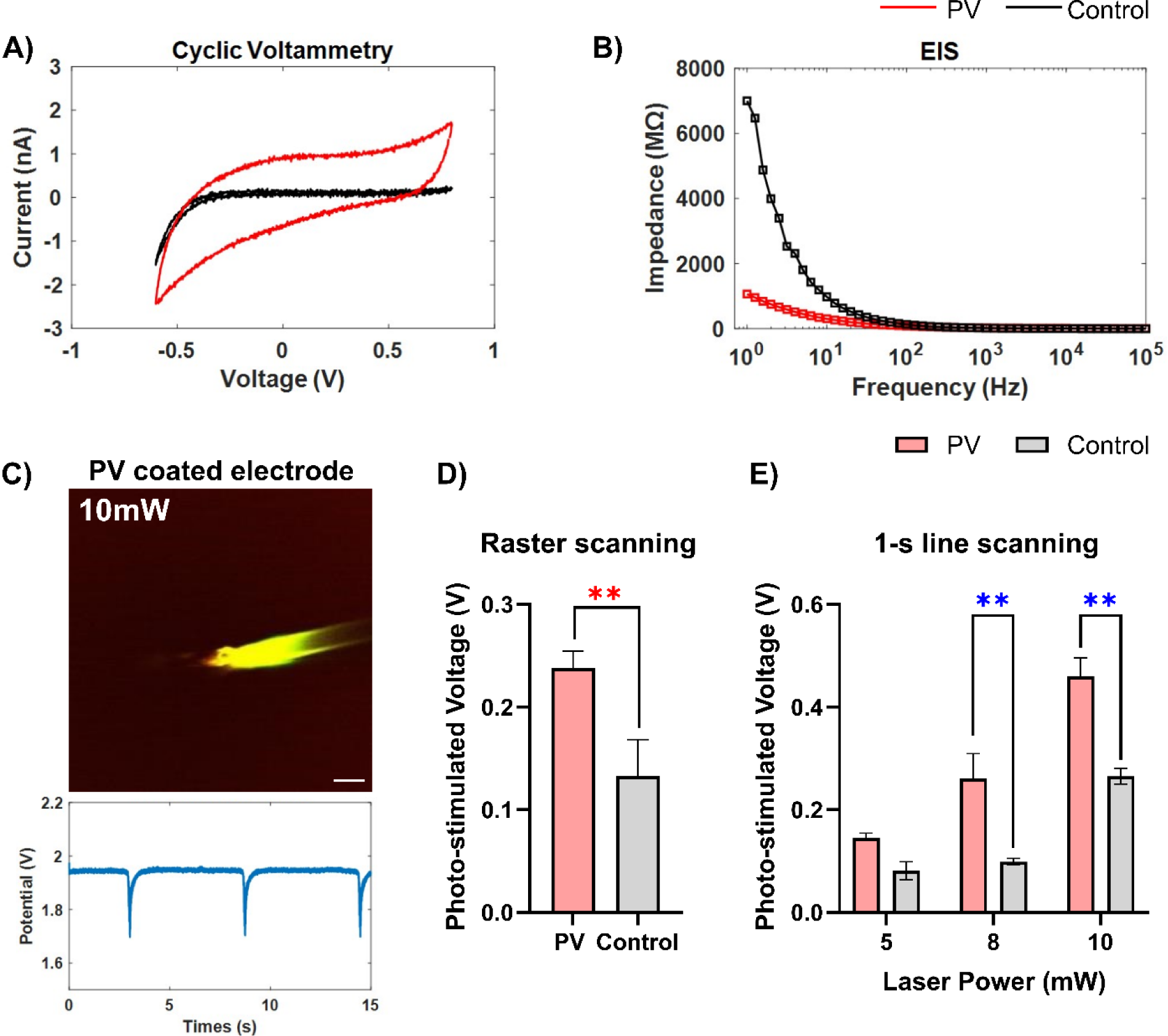
Photovoltaic (PV) polymer significantly increases photo-stimulated voltage at low laser power. A) Cyclic Voltammetry and B) Electrochemical impedance spectrum (EIS) of PV coated (red) and control (black) prototype electrodes. PV coating increased the charge storage capacity of the electrode. Additionally, PV coated electrode decreased the impedance, especially over the low frequencies. C) PV-coated electrode remained stable during 10 mW two-photon laser raster scanning and generated approximately 220 mV. Scale bar = 10 μm. D) PV coated electrode generated a significantly higher photo-stimulated voltage compared to uncoated controls by 10 mW raster scanning. Red double asterisks indicate the significant difference determined by Welch’s unpaired t-test, *p* < 0.05. E). PV coated electrode significantly enhances photo-stimulated voltage at 8 mW and 10 mW compared to uncoated controls. Blue double asterisks indicate the significant difference determined by Two-way ANOVA with Bonferroni corrected t-test, *p* < 0.005.

During the 10 mW laser raster scanning, parylene insulation of PV coated electrode was stable. The chronopotentiometry of PV coated electrode measured approximately 220 mV (**Fig. 4C**). Voltage generated by 10 mW rastering photo-stimulation at 800 nm wavelength, which was significantly increased compared to uncoated controls (**Fig. 4D**, unpaired t-test, p = 0.0018). Moreover, we examined the photo-stimulated voltages generated by the PV coated and the control electrodes during high duty-cycle scanning at different laser powers. Both PV coated and control electrodes increased photo-stimulated voltage in a laser-power dependent manner, suggesting carbon fiber electrodes are not damaged during this low power stimulation. There was no statistical difference in voltage between PV coated and control electrodes at 5 mW repetitive line scanning. However, PV coated electrode significantly enhanced the voltage at 8 and 10 mW laser power relative to uncoated controls (**Fig. 4E**, Two-way ANOVA with Bonferroni corrected t-test, p = 0.0221). In summary, the photovoltaic P3HT: PCBM polymer interfaces allow significant increases in photo-stimulated voltage.

It is important to note that even with the integration of photovoltaic polymers, carbon-based photoelectric devices may exhibit a relatively low efficiency in terms of neural stimulation. However, this type of photoelectric stimulation offers distinct advantages, particularly in achieving more precise neuronal activation. Unlike traditional tethered, current-based electrical stimulation that often results in a sparse distribution of activated neurons, photoelectric stimulation enables the activation of a more focal and discrete population of neurons [45, 64]. This targeted approach of driving nearby somas instead of passing axons helps reduce the risk of activating unintended neuron populations within neural circuits. Although several remote or wireless electrical stimulation devices such as the Floating Light-Activated Microelectrical Stimulator (FLAMES) have been proposed, these devices rely on larger stimulators compared to the ultrasmall photoelectric electrode implanted into the tissue, which could preferentially drive activation of passing axons and induce a larger immune response [30, 65, 66]. Additionally, a photoelectric electrode activated by NIR light may have a deeper penetration depth and a lower thermal gradient compared to the infrared light stimulation technique, due to its relatively shorter laser wavelength [67]. Moreover, in contrast to magnetic stimulation which is constrained by the bulkiness of coils and the extensive spread of magnetic fields [9], carbon-based photoelectric devices with ultrasmall size make them suitable for a wide range of applications such as in-field behavioral studies and dynamic neuroscience research. Thus, enhancing the efficiency of light-electric conversion, such as by applying photovoltaic polymers, helps to improve the potential of photoelectric devices in advancing our understanding and manipulation of neural activity.

### 3.4. *In Vivo* Evaluation of Prototype Electrode

To assess whether the strategies we previously discussed for the ultrasmall carbon electrode could improve the modulation of neural activity *in vivo*, we conducted photoelectric stimulation experiments using PV-coated and uncoated electrodes implanted in the Layer 2/3 of the visual cortex of Thy1-GCaMP6s mice. 1-s laser line scanning at 10 mW, 800 nm wavelength resulted in the activation of neurons in close proximity to the probe tip (**Fig. 5A-B**). Activated neurons of photo-stimulation were counted in binned distance away from the electrode. A majority of the activated neurons were located within a 50 μm radius from the electrode (**Fig. 5C**). The PV coated prototype electrodes resulted in a significantly larger number of activated neurons compared to the uncoated controls. These findings validated that the PV coating on the ultrasmall carbon electrode facilitates neuronal recruitment induced by photoelectric stimulation, paving the way for its potential future applications in neural modulation.

**Figure 5.**
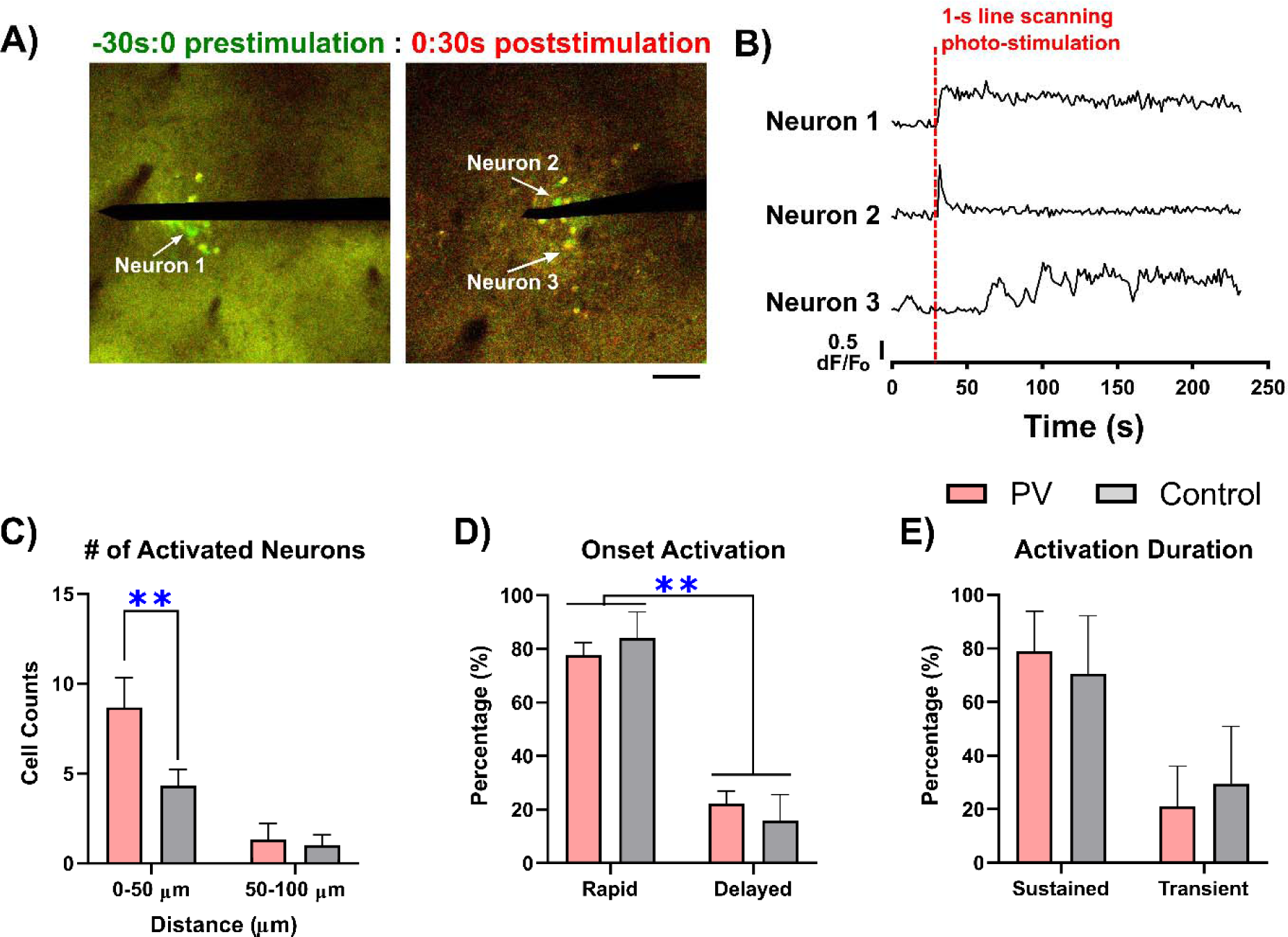
PV coated prototype electrode enhances neuronal recruitment in response to photo-stimulation. A) Merged images of the standard deviation projection over 30 seconds prestimulation (red) and poststimulation (green) in response to PV coated prototype electrode. Green neuronal soma denotes neuronal activation by 1-s line scanning photo-stimulation. Yellow indicates pixels before and after overlap. Prototype electrode was marked in black. Scale = 50 µm. B). Representative calcium traces show different profiles of neuronal activation in response to 1-s line scanning photo-stimulation. C) The counts of activated neurons by PV coated and uncoated control prototype electrodes at 0-50 μm and 50-100 μm distance. PV coated electrodes significantly increased the population of recruited neurons within 50 μm. D) The percentage of activated neurons with rapid onset was significantly higher than those with delayed onset by photo-stimulation of both PV coated and uncoated electrode. E). The percentage of activated neurons with sustained calcium elevation (> 30 s) and transient calcium activity (persisted < 10 s) by prototype electrode photo-stimulation. Blue double asterisks indicate the significant difference determined by Two-way ANOVA with Bonferroni corrected t-test, *p* < 0.05.

Interestingly, photo-stimulation tends to preferentially activate neuronal somas (**Fig. 5A**) rather than passing axons, a characteristic distinct from the typical scenario observed in faradaic electrical stimulation[46, 59, 68-74]. Previous computational studies have shown that the capacitive current at the soma and neurites are substantially different when compared to the sodium or potassium current [75]. Similarly, modeling studies have suggested that differences in the leading charge and waveform shape play a role in preferentially shifting excitation towards somas versus axons [74]. Selective activation is likely due to the differential impact of waveform shape on the charging or discharging of membrane capacitance in large somas and small axons before depolarization. This discrepancy can arise from the distinct membrane surface-to-intracellular volume ratios inherent in large somas and small axons. Experimental evidence supports the idea that waveform modulation can tune this preferential activation [68, 69]. However, it is crucial to note that electrical stimulation on IrOx electrodes, while demonstrating some preference modulation, still predominantly induces reversible faradaic charge injection due to the continuous application of electron current. Consequently, electrical stimulation significantly drives axonal stimulation. The capability of purely capacitive or photocapacitive stimulation to selectively activate local somas introduces novel possibilities for manipulating neuronal networks through stimulation strategies.

Here we observed that neurons activated by photo-stimulation displayed diverse calcium dynamics over a 200-second post-photo-stimulation period. The representative calcium profiles of these neurons revealed distinct patterns in both the onset and duration of activation (**Fig. 5B**). The PV-coated electrode resulted in 77.69 ± 6.51% of activated neurons showing rapid onset of calcium elevation, which is comparable to the 79.37 ± 8.98% observed with the uncoated control (**Fig. 5D**). Additionally, the percentage of activated neurons with sustained calcium activation was comparable between the PV coated and control prototype electrode (**Fig. 5E**, PV coated:78.89 ± 21.14%, uncoated control: 70.63 ± 30.51%). Together, these results provide a characterization of neuronal recruitment patterns in photo-stimulation of prototype electrodes, showing local neuron activation by photoelectric stimulation primarily exhibits rapid onset and sustained activation. It was further observed that the PV coating does not significantly alter this foundational response pattern.

Organic photovoltaic polymers have been applied in extracellular stimulation of neurons and thus have great potential for *in vivo* biological applications, such as retinal prostheses [50]. In-vitro studies demonstrate that these photovoltaic polymer interfaces can convert light entering the pupil into electrical stimuli, serving as artificial photoreceptors to activate retinal cell [50, 53, 76]. Our findings indicate that a photovoltaic P3HT:PCBM polymer coating enhances voltage generation and neuronal activation selectivity in the Thy1-GCaMP visual cortex (Fig. 5), confirming its potential for *in vivo* cortical neuron modulation. However, the mechanism of P3HT:PCBM blends in a biological environment may differ from those in solar cells where bulk heterojunctions facilitate the charge pair separation and limit the charge recombination [48]. In contrast, it appears that photoelectric-based neuromodulation is driven more by capacitive charging at the polymer-electrolyte interface rather than direct electron transfer [51]. Consequently, the electron donor-acceptor interface may not be the most effective for neuronal activation. Studies suggest that the donor component only (P3HT) can efficiently stimulate cultured neurons when illuminated [50, 77]. Future research should explore the impact of using P3HT-only coating on photo-stimulated voltage and its potential for *in vivo* neuronal stimulation.

### 3.5. Potential of Ultrasmall Carbon-Based Diamond Electrodes for Photo-Stimulation

While our prototype electrode that utilizes a carbon fiber core and parylene insulation could successfully stimulate neural activity, the limitation was recognized regarding the stability under high laser intensity stimulation. Therefore, we briefly investigated the potential of alternative conductive material derived from carbon compounds on photoelectric stimulation. Diamond is a form of carbon where it is sp3 bonded and has a robust and rigid bonding structure. When highly doped with boron, it creates defect sights, which allow diamond to be conductive [78–81]. Diamond has a bandgap of 2.34 eV, and as the diamond is further doped, the band gap is tuned and reduced, as the material becomes more and more metallic like [82]. Here we tested a BDDME, where the BDD is insulated in a glass capillary. This BDD has demonstrated several advantages, including an exceptionally wide potential window, low background current, and good biocompatibility [78]. Thus, the diamond electrode has the potential to be applied in neurochemical detection with high spatiotemporal resolution and sensitivity.

Here, we ask the question of whether the diamond electrode can generate photo-stimulated voltage to safely stimulate neural tissue under multiphoton laser scanning. Two photon time series images demonstrate the integrity of the diamond electrode by 40 mW photo-stimulation. Quantification of the Thy1-GCaMP cell density revealed a significant local neuronal activation near the photo-stimulated diamond electrode (**Fig. 6**, One-way ANOVA with Bonferroni corrected t-test, p = 0.0076). In summary, a diamond electrode as an alternative carbon-based ultrasmall electrode can lead to a sustained increase in neural activity near the implant.

**Figure 6.**
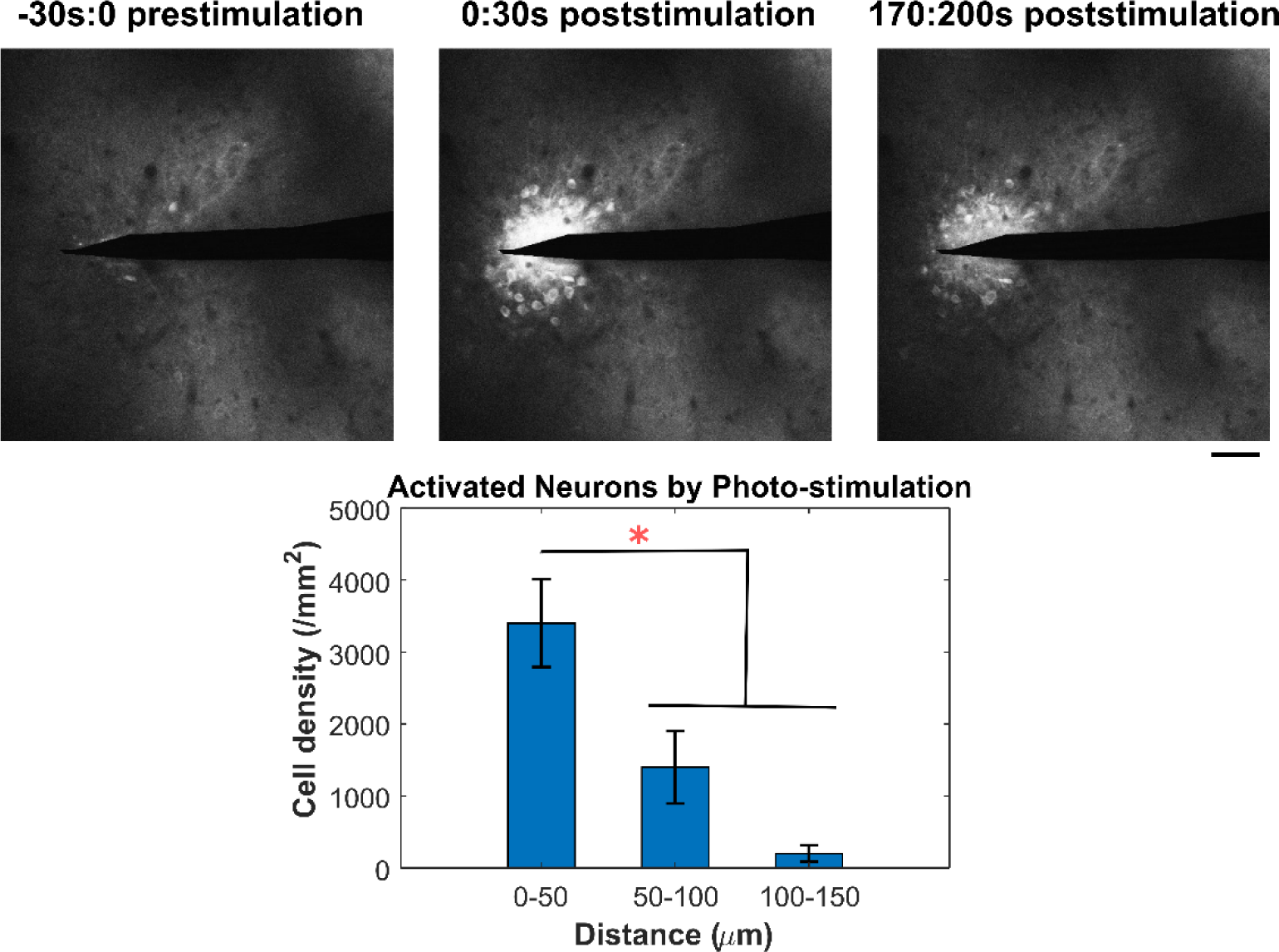
Boron doped diamond electrode successfully led to local neuron activation at 40 mW photo-stimulation. Photostimulation leads to significant increases in the cell density of activated neurons within 50 µm relative to 50-150 µm distance away from the stimulated diamond electrode. Scale bar = 50 µm. Red double asterisks indicate the significant difference determined by One-way ANOVA with Bonferroni corrected t-test, *p* < 0.05.

Diamond offers a promising alternative material for photoelectric-based neuromodulation, due to its tunable bandgap, and high biocompatibility. During diamond growth and fabrication, it can be grown intrinsically, with varying grain sizes, or it can be doped with boron, to enable high metallic like conductivity. Insulating it in glass, and epoxy, the transparency allows light energy to reach the conductive, boron-doped diamond for subsequent light-electric conversion, showcasing this phenomenon, to our knowledge, the first reported instance. During high-power multiphoton stimulation, *in vivo* observations showed no noticeable damage or formation of gas bubbles (**Fig. 6**). This is likely attributable to the reduced sp^2^ carbon content of the diamond and the robust mechanical properties, and which enable it to maintain its integrity even when subjected to the disruptive influence of stimulation light energy. Future next steps are to encase the conductive diamond in intrinsic polycrystalline diamond (PCD) cladding, where the all-diamond electrode has good biocompatibility and serves as an effective dielectric barrier to prevent signal cross-talk [78, 79]. Moreover, methods for processing and patterning diamond enable high-throughput batch fabrication and customization of electrode arrays with unique architectures [78]. This procedure not only reduces manufacturing costs but also enhances design flexibility [78, 83, 84]. Additionally, the performance of diamond electrodes can be finely tuned by manipulating the boron doping level. Highly boron-doped diamond enhances sensitivity and electron transfer kinetics and thus makes it suited for neurotransmitter detection applications [78]. Together, diamond could have promising applications in untether, ultrasmall photoelectric-based stimulation devices. Future studies should focus on optimizing device design and processing parameters to reveal their full potential.

### 3.6. Precise Drug Delivery by Photo-Stimulation of Ultrasmall Carbon Fiber Electrode

As untethered, ultrasmall photoelectric devices possess the capability to electrically modulate neural tissue, we then ask the question whether these devices have the potential for biochemical modulation. One strategy involves the integration of a conductive polymer with dopants at the tissue-electrode interfaces. These conductive polymers can have their chemical and electrical properties easily engineered to enable electrically controlled drug delivery of loaded neurochemicals and biomolecules [16, 85–88]. Dopants such as SNP as porous nanoscale carriers help overcome the limited surface of carbon-based photoelectric electrodes, which enhance drug loading capacity and allow for fine-tuning of the release rate [89–91]. Therefore, we investigated the potential of the Poly(3,4-ethylenedioxythiophene) (PEDOT) conductive polymer doped with SNP for achieving controlled drug release by photo-stimulated carbon fiber electrodes.

To quantify the drug release capabilities of this coating, we employed fluorescein as a model drug. Fluorescein was loaded into SNPs which are incorporated into PEDOT through an electrochemical deposition process on carbon fiber electrodes. The release and dispersal of the fluorescein dye were evaluated in agarose gel. We observed that the 0.2 Hz rastering scanning resulted in a substantial release of the fluorescein dye (**Fig. 7A**).

**Figure 7.**
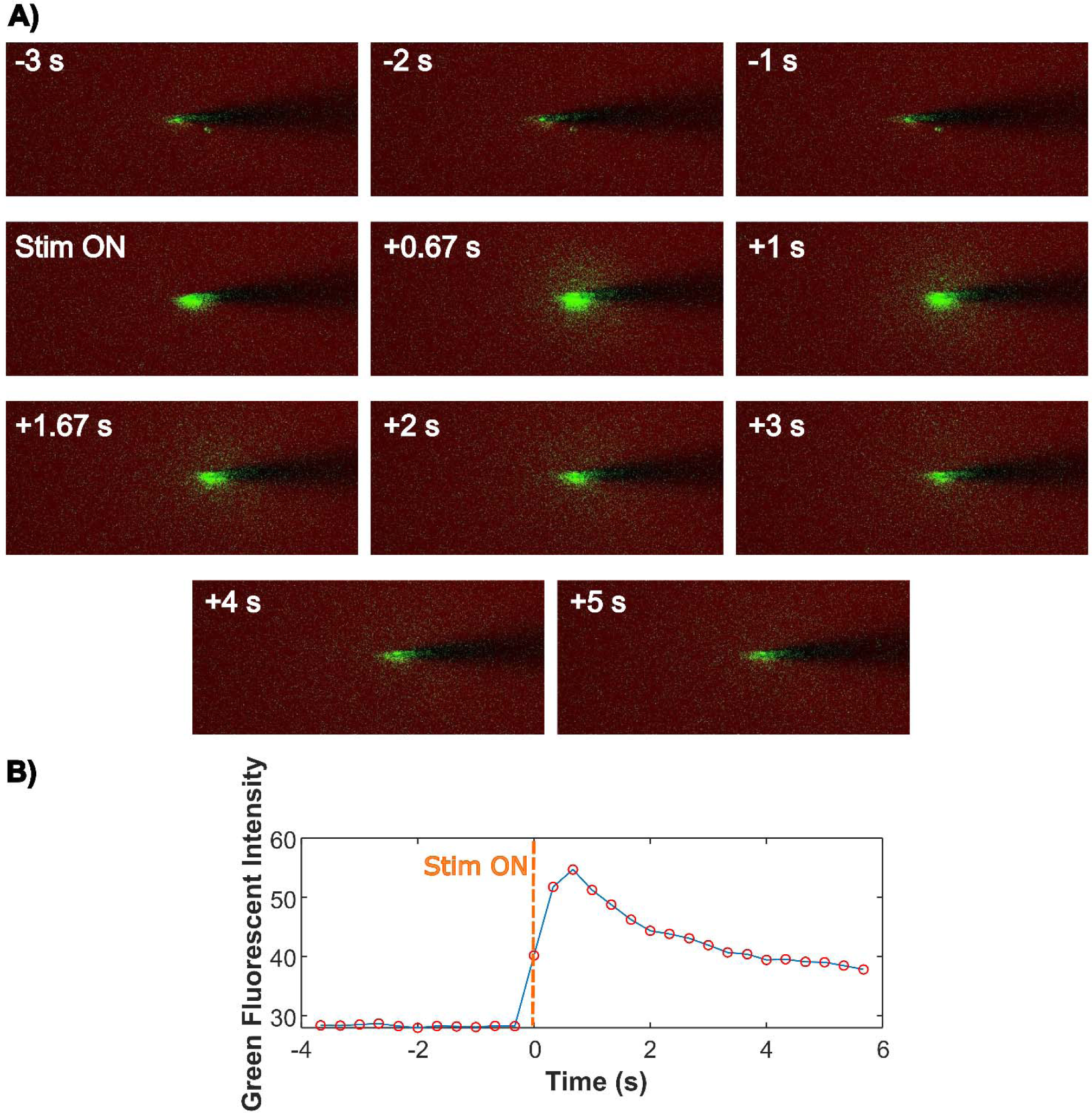
Ultrasmall carbon fiber electrode with PEDOT/SNP coating releases loaded fluorescein at 10 mW photo-stimulation. A) A) Representative time series images of the green fluorescein before, during, and after photo-stimulation. Scale bar = 10 µm. B) Quantification of the green fluorescent intensity of the fluorescein-loaded carbon fiber electrode over time.

Once released, the dye rapidly diffused into the surrounding agarose gel within several seconds (**Fig. 7B**). The temporal release profile of the fluorescein dye shows that photostimulation of the PEDOT/SNP/fluorescein coated ultrasmall carbon fiber electrode can lead to localized and controllable release of the loaded drug. These findings demonstrated that combination of doped conductive polymer coating with photoelectric electrodes is promising for on-demand drug delivery, which offers a unique capability wherein light can precisely control the release of biomolecules with both temporal and spatial accuracy (Fig. 7). The PEDOT/SNP system presents a versatile platform for loading and releasing a wide range of drugs, including essential neurotransmitters such as glutamate, the inhibitory transmitter GABA, as well as dopamine (DA), 6,7-Dinitroquinoxaline-2,3-dione (DNQX), and bicuculline [89, 90, 92]. Thus, it holds promise for biochemical modulation using light-controlled, minimally invasive photoelectric electrodes to study the mechanisms of neural circuits involved in both normal and pathological cognition. This approach enables the focused modulation of neuronal activity in untethered, freely moving animals. However, there exist several challenges that hinge on the widespread application of this combined technology in neuroscience research. Firstly, it remains uncertain whether the loaded drug reservoir is sufficient for sustaining long-term release for chronic neuromodulation. Secondly, the thickness for PEDOT/SNP coating should be optimized for drug release performance, striking a delicate balance between the requirement of sufficient drug supply and the inherently small dimensions of the photoelectric electrode. Lastly, the stability and integrity of the PEDOT/SNP coating on the photoelectric electrode during photo-stimulation should be evaluated to address potential concerns related to biocompatibility and electrode functionality. Future research endeavors aimed at improving the integration of photoelectric stimulation technology and doped conductive polymer coatings should consider these challenges.

## 4. Conclusion

The neuromodulation technologies that are crucial for treating neurological disorders and exploring basic neuroscience are rapidly developing. Stimulation based on the photoelectric effect is a novel tether-free strategy to electrically modulate brain tissue by external light sources. This stimulation strategy allows the carbon fiber electrode to generate non-Faradic, capacitive charge transfer at the electrode-electrolyte interface, and the ultrasmall dimension of the carbon electrode also reduces the degree of brain tissue injury. To improve the potential of this photoelectric stimulation with ultrasmall carbon electrode for wider application, we investigated diverse strategies including the flexible insulation material, laser stimulation strategy, photovoltaic polymer interface coating, alternative carbon-based doped diamond electrode, and integration with drug delivery system. Our findings indicate that flexible and transparent parylene-C insulation is unsuitable for high-intensity laser photo-stimulation due to susceptibility to damage. We found that strategies of high duty-cycle laser stimulation and photovoltaic polymer coating enhance the voltage of the cathodic pulse generated by multiphoton laser stimulation. Boron-doped diamond electrode results in sustained calcium activation in local neurons. Moreover, the photo-stimulation of alternative carbon-based diamond electrodes demonstrated sustained neuronal activation *in vivo* while preserving electrode integrity even at high laser intensities. Additionally, our study highlights the potential of integrating photoelectric stimulation with PEDOT/SNP conductive polymer coating for on-demand drug delivery. These results several promising avenues for enhancing photoelectric stimulation technology, which will help expand the horizons of photoelectric stimulation development in both research and clinical applications.

## Acknowledgements

This work was supported by NIH R01NS105691, and NSF CAREER CBET 1943906 to Kozai and NIH R01NS110564-01 and NSF 1926756 to Cui.

## Credit

**Keying Chen**: Conceptualization, Methodology, Validation, Project administration, Investigation, Data Analysis, Data curation, Writing – original draft, review & editing. **Bingchen Wu**: Methodology, Validation, Investigation, Writing – review & editing. **Daniela Krahe**: Methodology, Validation. **James Siegenthaler**: Methodology, Electrode fabrication. **Robert Rechenberg**: diamond fabrication and structuring. **Wen Li**: Conceptualization, Methodology. **Tracy Cui**: Conceptualization, Supervision. **Takashi D.Y.Kozai** : Conceptualization, Methodology, Resources, Writing – review & editing, Supervision, Funding acquisition.

